# Natural genetic variation in *C. elegans* reveals genomic loci controlling metabolite levels

**DOI:** 10.1101/217729

**Authors:** Arwen W. Gao, Mark G. Sterken, Jelmi uit de Bos, Jelle van Creij, Rashmi Kamble, Basten L. Snoek, Jan E. Kammenga, Riekelt H. Houtkooper

**Affiliations:** Laboratory Genetic Metabolic Diseases, Academic Medical Center, 1105 AZ Amsterdam, The Netherlands; Laboratory of Nematology, Wageningen University and Research, 6708 PB, Wageningen, The Netherlands.

## Abstract

Metabolic homeostasis is sustained by complex biological networks responding to nutrient availability. Disruption of this equilibrium involving intricate interactions between genetic and environmental factors can lead to metabolic disorders, including obesity and type 2 diabetes. To identify the genetic factors controlling metabolism, we applied a quantitative genetic strategy using a *Caenorhabditis elegans* population consisting of 199 recombinant inbred lines (RILs) originally derived from crossing parental strains Bristol N2 and Hawaii CB4856. We focused on the genetic factors that control metabolite levels and measured fatty acid (FA) and amino acid (AA) composition in the 199 RILs using targeted metabolomics. For both FA and AA profiles, we observed large variation in metabolite levels with 32-82% heritability between the RILs. We performed metabolite-metabolite correlation analysis and detected strongly co-correlated metabolite clusters. To identify natural genetic variants responsible for the observed metabolite variations, we performed QTL mapping and detected 36 significant metabolite QTL (mQTL). We focused on the mQTL that displayed high significant linkage and heritability, including an mQTL for the FA C14:1 on chromosome I, and another mQTL for the FA C18:2 on chromosome IV. Using introgression lines (ILs) we were able to narrow down both mQTL to a 1.4 Mbp and a 3.6 Mbp region, respectively. Overall, this systems approach provides us with a powerful platform to study the genetic basis of *C. elegans* metabolism. It also allows us to investigate additional interventions, such as nutrients and stresses that maintain or disturb the regulatory network controlling metabolic homeostasis, and identify gene-by-environment interactions.

## Introduction

Energy homeostasis is maintained by biological networks that respond to nutrient availability and keep functional balance at cellular and molecular levels (Andreux et al. 2012). When this equilibrium is disrupted by a combination of genetic and environmental factors, such an imbalanced metabolic system can lead to metabolic disorders, including obesity and type 2 diabetes (Andreux et al. 2012). To study the interactions between genetic and environmental factors, different approaches are used, including reverse and forward genetics (Williams and Auwerx 2015). Reverse genetics approaches comprise techniques that focus on the phenotypic impact of the knockdown, knockout, or overexpression of specific candidate genes. This approach typically focuses on a single gene, and therefore has several limitations (Williams and Auwerx 2015): (1) the additive and non-additive interactions between gene variants cannot be observed; (2) common variants with a subtle effect cannot be detected; (3) prior hypotheses about the gene function are a prerequisite. Instead, forward genetics bypasses these limitations as it exploits the natural phenotypical variation in a population to identify causal genetic variants (Williams and Auwerx 2015). This involves classic mutagenesis screens and techniques such as quantitative trait loci (QTL) analysis and genome-wide association studies (GWAS). QTL analysis is a statistical technique that examines the association between a marker genotype and a quantitative trait, *i.e.* a trait with continuous phenotypic variation that is affected by genetic and/or environmental factors. For QTL analysis, a segregated population is used to find genomic regions that are associated with the trait variation in the population (Slate 2005). In contrast, GWAS relies on natural populations to identify common genetic variants associated with traits, and has been successful in identifying many loci associated with susceptibility to complex diseases such as type 1 and type 2 diabetes (Visscher et al. 2017). In this study, we chose to use the nematode *Caenorhabditis elegans* as a model organism to explore genetic variation affecting metabolic parameters. Considerable variation would validate the nematode for further identification of complex gene by environment interactions correlated with metabolism.

*C. elegans* is a versatile model organism for understanding complex genetic pathways underlying distinct phenotypes such as stress response, lifespan, host-pathogen interaction, and behaviors (Gao et al. 2017b). The Kammenga group generated a segregating population of *C. elegans* derived from the genetically and ecologically divergent strains N2 (Bristol) and CB4856 (Hawaii), that is suitable for studying genome-to-phenome relations (Li et al. 2006; Andersen et al. 2012; Sterken et al. 2015). The segregating population consists of 199 homozygous recombinant inbred lines (RILs) that have been SNP-genotyped and phenotyped for several traits, including gene expression (Li et al. 2006; Rockman et al. 2010; Vinuela et al. 2010) and stress-response hormesis (Gutteling et al. 2007; Doroszuk et al. 2009; Rodriguez et al. 2012; Snoek et al. 2017). The 199 RILs all have a different combination of alleles derived from either the N2 or CB4856 parental strains, allowing us to determine the natural influence of genetics on a phenotypic trait.

In *C. elegans*, changes in metabolism are often studied at the transcriptional level, and indeed multiple metabolic aging genes have been identified in worms (Houtkooper et al. 2013; Lopez-Otin et al. 2013; Gao et al. 2017b). At the same time, to get a comprehensive understanding of the metabolic changes underlying the aging process, one has to get closer to measuring metabolic physiology, for instance using metabolomics. Based on recent advances in the metabolomics field, our group developed a sensitive mass spectrometry (MS)-based platform for measuring metabolites in *C. elegans*. This platform allows us to detect 44 medium-chain, long-chain, and very-long-chain fatty acids (FAs) (C14:0-C30:0) and 19 amino acids (AAs) in a sample of around 500 worms (Gao et al. 2017a). Using this platform, we measured metabolites in the RILs of *C. elegans* on a large-scale, and identified the effect of genetic variation on metabolism (Figure 1). We measured the FA and AA profile of a panel of 199 RILs to identify the variable genomic loci affecting metabolite levels. Overall, we systematically investigated the genetic basis of metabolism by QTL mapping showing that this RIL panel is a powerful platform to study complex metabolic traits that underlie interactions between genes and environmental factors.

**Figure 1.**
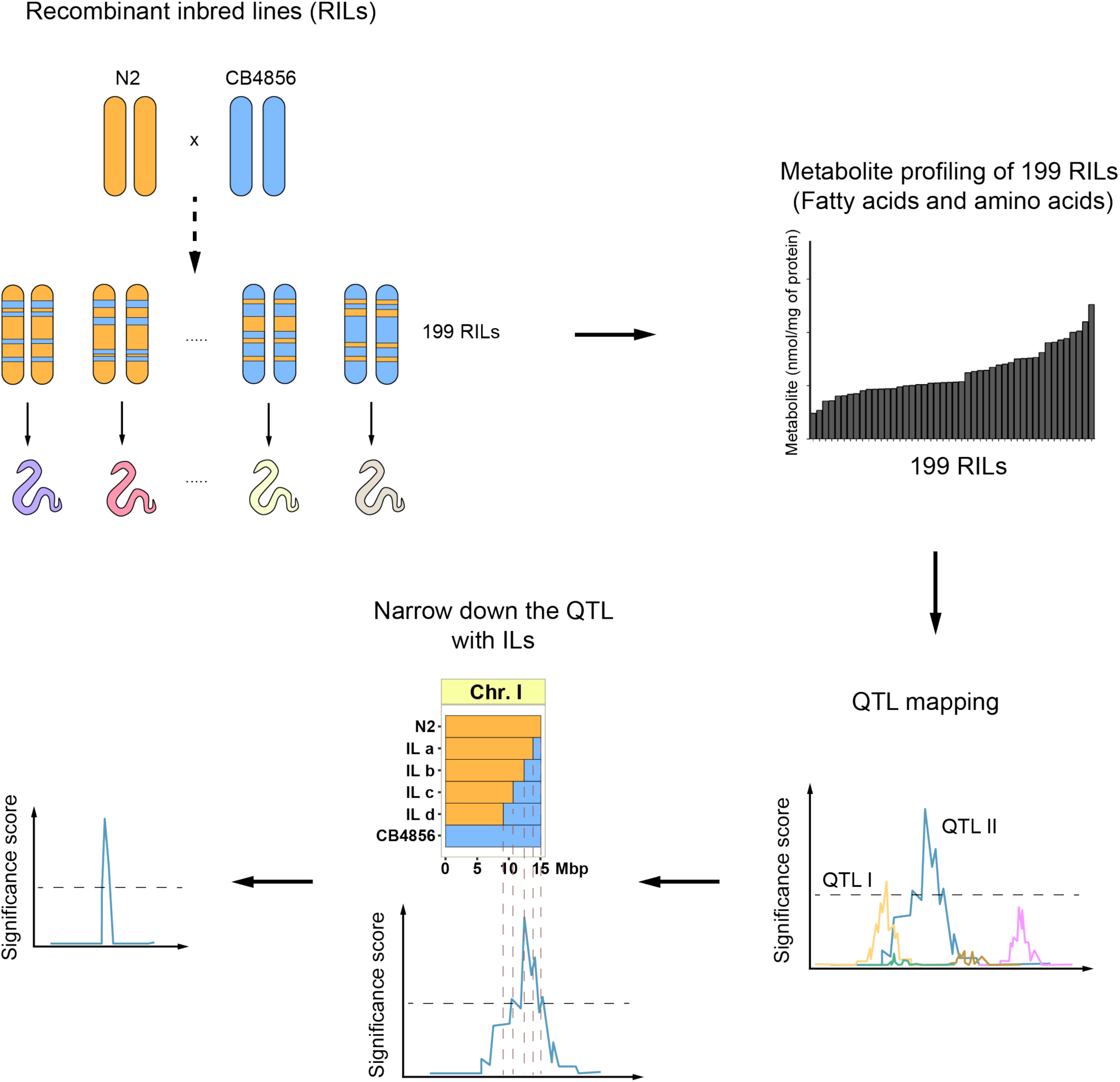
Flowchart of the strategy to identify genetic loci that control metabolic traits. Metabolite profiles for 19 amino acids (AAs) and 44 fatty acids (FAs) were measured in 199 recombinant inbred lines (RILs). These metabolite profiles were plotted and correlated with the RIL genetic map, identifying genetic loci linked to variation in metabolite levels (metabolite quantitative trait loci, mQTL). Subsequently, these mQTL were verified/confirmed and fine-mapped using introgression lines (ILs), strains containing a small locus of one parental strain in the genetic background of the other parental strain (N2: orange; CB4856: blue).

## Results

### Metabolite levels in C. elegans show high heritable variation

To extract general principles about the complex genetics of metabolism, we measured the impact of genetic variation on metabolite levels in 199 recombinant inbred lines (RILs) (Supplementary file 1) using our recently established sensitive MS platform (Figure 1) (Li et al. 2006; Thompson et al. 2015; Gao et al. 2017a). The metabolite traits in the RILs displayed high levels of variation between the RILs, sometimes beyond what was observed in the two parental strains, which suggests transgression (or transgressive segregation) (Figure 2A-B). For instance, the most abundant metabolites, C18:1 and alanine, showed a large difference between the lowest and highest line of 23.0- and 68.9-fold, respectively (Figure 2B). Also, the metabolites present in nematodes at lower concentrations, including C20:3 and methionine, showed a large difference between the lowest and highest RIL of 59.0 and 32.1-fold, respectively. Transgression was observed for 18 metabolites, especially in FAs (FDR = 0.05; Supplementary file 3). Transgression analysis is a measure for the genetic complexity of a trait, for example due to genetic interactions. It showcases that metabolite levels are affected by multiple polymorphic genes of which the alleles cause a balanced phenotype in the parental lines.

**Figure 2.**
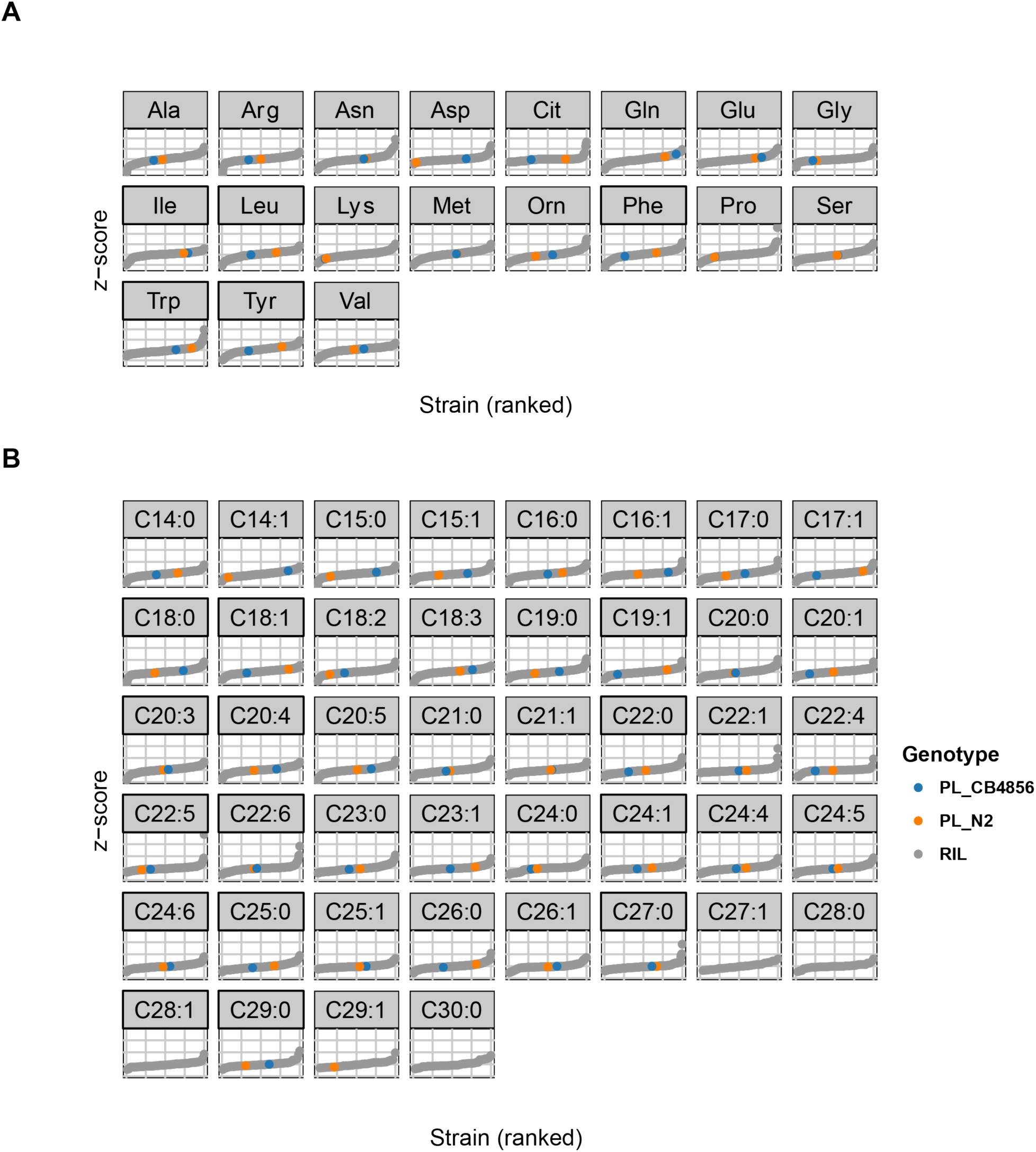
Metabolite levels across the 199 RILs and the two parental strains. (A) The average trait level of 19 AAs was first expressed as percentage of total AAs, followed by *z*-score transformation. The 199 RIL values were indicated in grey, the value of the parental strain N2 in orange, and the value of the parental strain CB4856 in blue. Trait levels below limit of quantification of AA measurement (0.4 nmol/mg of protein) were not shown. (B) The mean values of the level of 44 FAs were shown after data transformation (the absolute FA concentration was expressed as a percentage over total FAs, followed by z*-* score transformation). The FA levels in 199 RILs were indicated in grey, that in the parental strain N2 and CB4856 were indicated in orange and blue, respectively. FA levels below limit of quantification of the measurement (0.03 nmol/mg of protein) were not shown.

Next, we calculated the broad-sense heritability (H^2^) of these metabolite traits, which is a measure for the genetic contribution to the variation in metabolite levels (Figure 3). For more heritable traits, the contributing genomic loci (quantitative trait loci, QTL) are more likely to be found. We found significant heritability for 51 metabolite traits (FDR = 0.05), ranging from 0.32 (tryptophan) to 0.69 (lysine) for the AAs (Figure 3A), and from 0.32 (C22:1) to 0.82 (C14:1) for the FAs (Figure 3B; Supplementary file 4). Together, our data show that both AAs and FAs have moderate to high heritability, making QTL identification likely. Specifically, the traits with non-significant transgression and high heritability, for instance the FA C14:1 (H^2^ = 0.82), are most likely associated to a single major locus explaining most of the metabolic variation.

**Figure 3.**
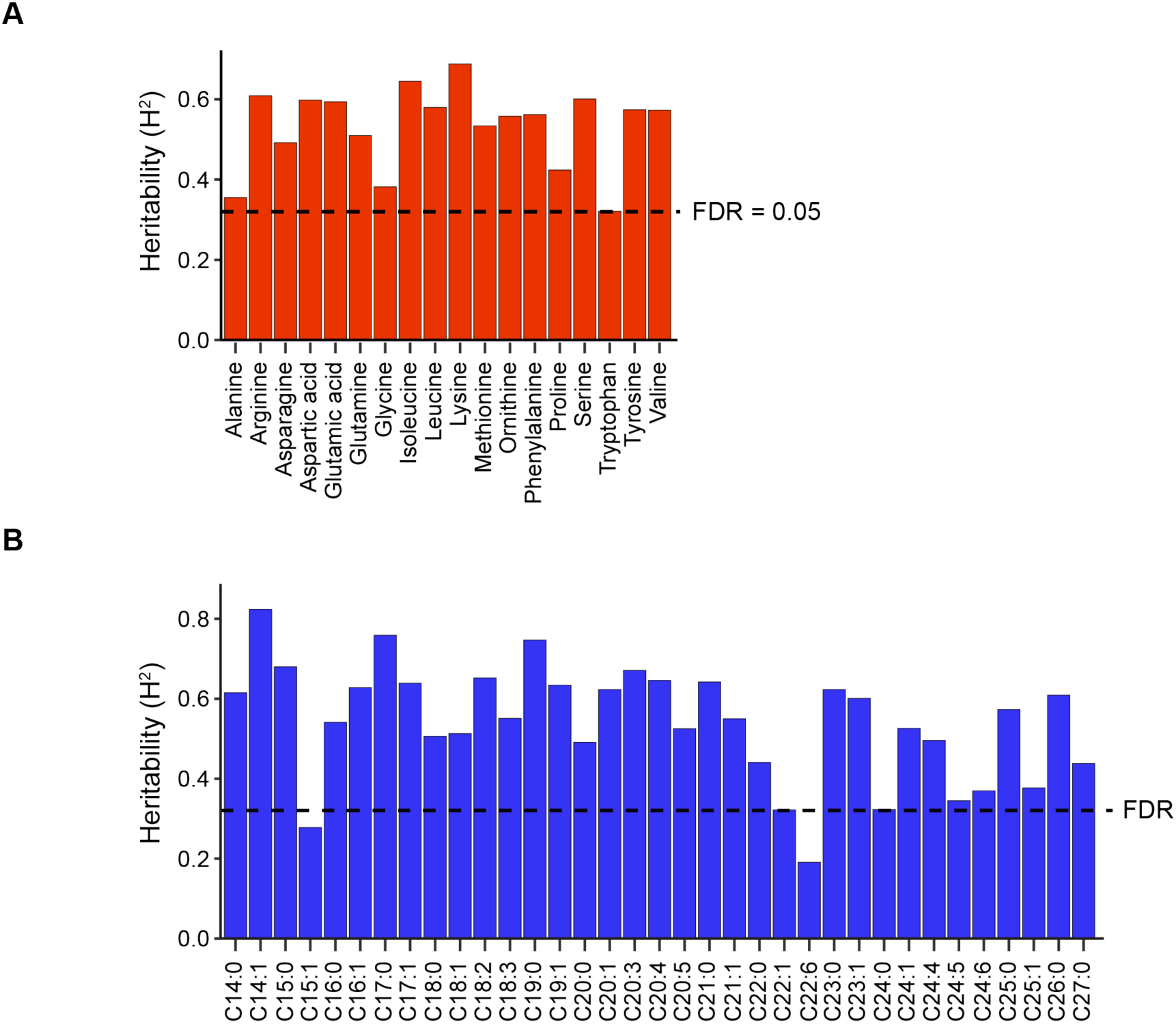
Heritability analysis of AAs and FAs. We estimated broad-sense heritability (H^2^) of metabolic traits based on a set of 51 RILs that were measured at least three times. The bars indicate the H^2^, where the dashed line indicates a permutation-based false discovery rate (FDR) of 5% (based on 1,000 permutations). We found evidence for strong, significant, heritable variation in most AA (**A**) and FA (**B**) abundances, ranging from 0.32 – 0.82 (FDR = 0.05). Only for FA C15:1 and C22:6 no significant H^2^ was found.

### Metabolite levels exhibit strong correlations in the RIL cohort

Levels of different metabolites are likely correlated since homeostasis is supported by integrated metabolite networks (Houtkooper et al. 2010). To investigate this, we calculated the correlations for all pairs of FAs and AAs over all RILs independently. We found strong correlations between different clusters of metabolites (Figure 4). In the correlation analysis of AAs, we observed a strong positive correlation cluster between several hydrophobic AAs, including methionine and phenylalanine, the hydrophilic AA tyrosine, and branched-chain AAs (BCAAs) valine, leucine, and isoleucine (Figure 4A). Another positively correlated cluster was found in ornithine, citrulline, glycine, serine, lysine, glutamic acid and aspartic acid (Figure 4A). Interestingly, this cluster negatively correlated with the former positive cluster. Overall, we observed clear connections and correlations between many metabolites, suggesting shared genetic variants that regulate the levels of these metabolites.

**Figure 4.**
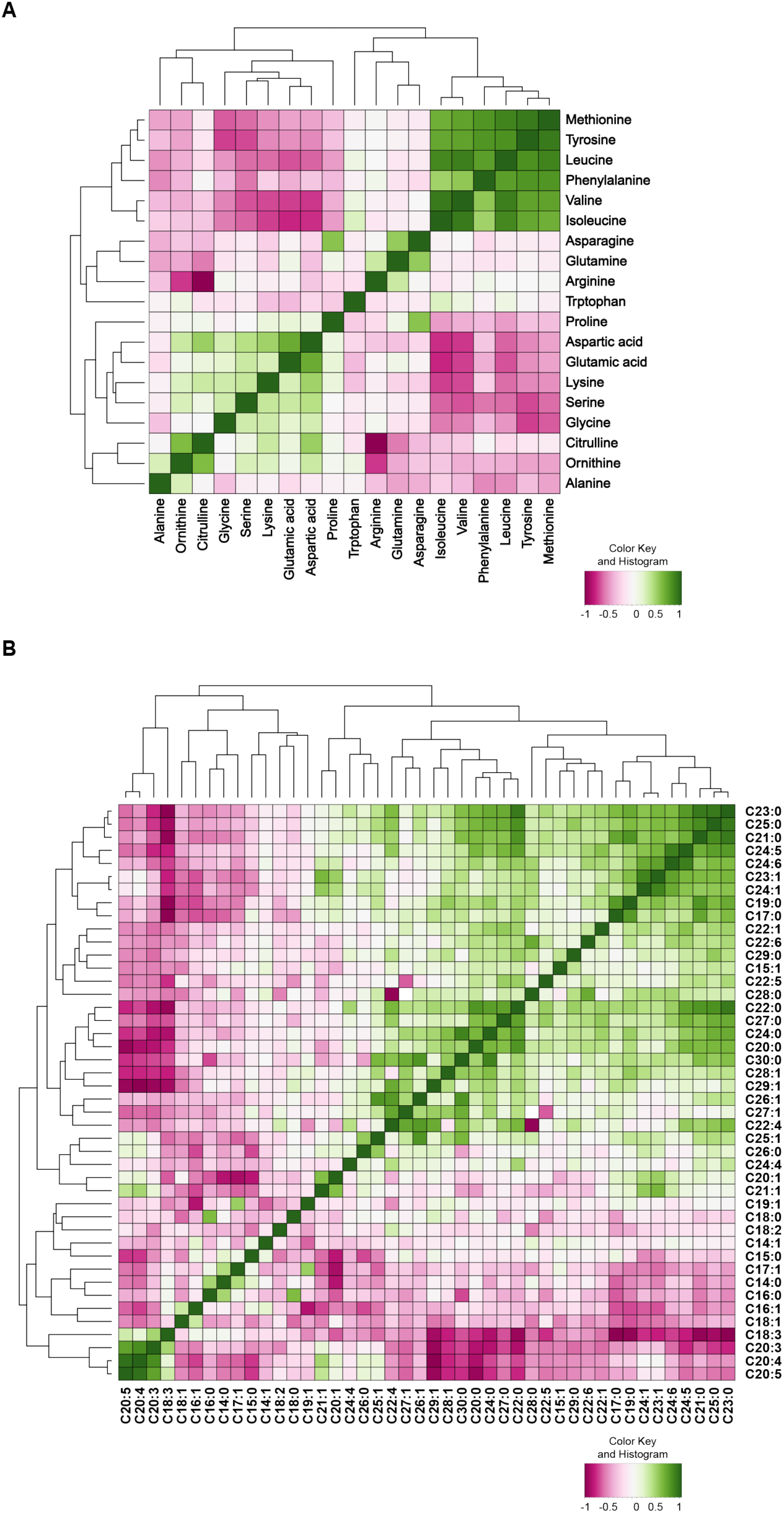
Correlation analysis of AA and FA species in the RIL strains. (A) Correlation heat map of AA profiles of all the RIL strains. There was a strong cluster of methionine, tyrosine, phenylalanine, arginine and branched-chain amino acids (BCAAs) leucine, isoleucine and valine. Another positive correlated cluster was found in ornithine, citrulline, glycine, serine, lysine, glutamic acid and aspartic acid. Interestingly, the former positive cluster was negatively correlated with some AAs from the later cluster, including proline, aspartic acid glutamic acid, lysine, serine, and glycine. (B) Correlation heat map of FA profiles of the RILs. Strong correlation was found a group of long chain and very long chain FAs. Polyunsaturated fatty acids (PUFAs) were also clustered and positively correlated.

In the FA correlation profile, we observed a strong positive correlation in several polyunsaturated FAs (PUFAs), including C18:3, C20:3, C20:4, and C20:5 forming a small cluster (Figure 4B). Strong correlations were also found in a large cluster of long-chain and very long-chain FAs. Another large cluster was formed by C30:0 and several unsaturated FAs (e.g C20:1, C22:1, C24:5, and C24:6) (Figure 4B). Notably, these last two clusters showed a strong negative correlation with the first small cluster. Taken together, the strong correlations within metabolite classes imply that linked metabolite ((m)QTL) could be detected as many metabolites show similar patterns of variation over the RILs.

### Multiple QTL link metabolite levels to causal loci

To identify loci that explain variation in metabolite levels, we performed QTL mapping on 56 metabolites measured in the 199 RILs (Figure 5A). We expected to detect 80% of the genomic loci explaining a minimum of 10% per locus of the trait variation in this population when using a single marker model when using the –log10(p) > 3.7 as the significance threshold (Supplementary file 6). We detected 36 significant mQTL for 26 metabolites (Figure 5A; Supplementary file 7). This shows that specific loci affecting metabolite trait variation can be identified.

**Figure 5.**
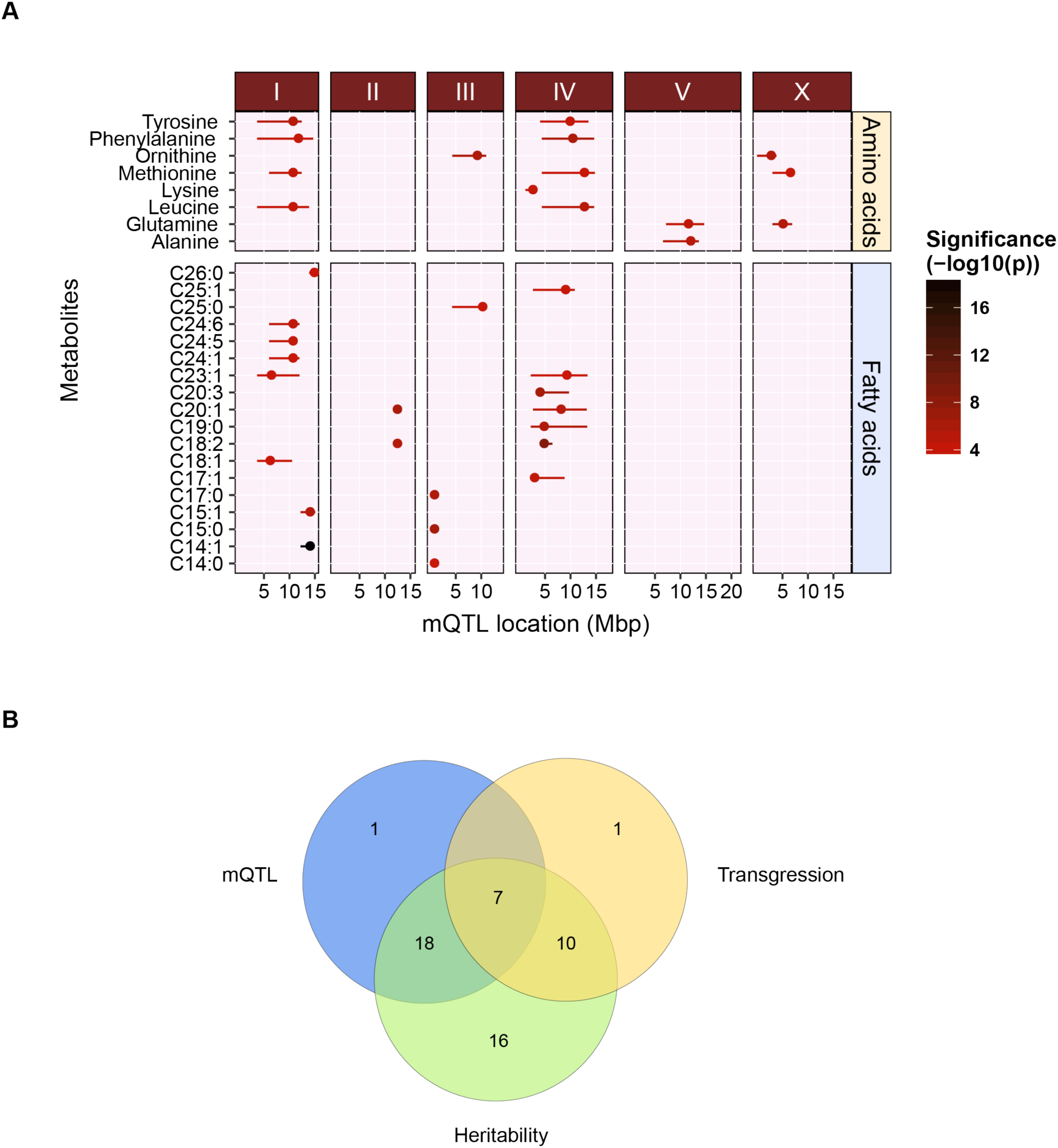
QTL mapping of metabolite levels in the 200 RILs. (A) To identify genetic factors responsible for the observed metabolite trait variations, we performed QTL mapping with the metabolite profiles of 199 RILs. In total, 36 significant mQTL were detected for AA and FA traits at an FDR = 0.05 (–log(p)>3.7) ranging from 4.2 to 16.5. The x-axis displayed the position of the QTL in mega base pairs (Mbp) for each chromosome and the y-axis displayed the trait for which a significant QTL was found (B) Overlapping metabolite traits that are identified in QTL mapping, heritability analysis and transgressive segregation analysis. Except for one metabolite trait (C15:1), all metabolite traits that were mapped to one or more significant QTL were highly heritable. Seven transgressive metabolite traits appeared to be associated with one or more significant QTL. These seven traits also have significantly higher heritability.

For the AA traits, we observed 15 significant mQTL for eight different AAs. We found that four AAs (tyrosine, phenylalanine, methionine, and leucine) shared two broad mQTL peaks on chromosome I (~10.5 Mbp) and chromosome IV (~11.0 Mbp). These four AAs also displayed strong positive correlations between the trait levels in the RILs (Figure 4A). For the AA methionine, an additional mQTL was found on chromosome X, bringing the total number of mQTL for this AA to three. Individually, each of these three QTL explained ~7% of the total variation and together they explained 18.5% of the variation in methionine levels between the RILs in an additive model. Furthermore, only two AAs, lysine and alanine, were each mapped to a single locus, on chromosome IV and V, respectively.

For the FA traits, we detected a total of 21 significant mQTL for 18 unique FAs. Among these FAs, we observed several shared mQTL. For instance, we detected a mQTL in the first 0.6 Mbp of chromosome II for C14:0, C15:0, and C17:0. Also, the mQTL for three very-long-chain FA species C24:1, C24:5, and C24:6 were all mapping to a locus on chromosome I, with a peak at 10.5 Mbp. Across the QTL analysis, the most significant mQTL we detected (-log10(p) = 18.8), was for C14:1, which mapped to a single locus on chromosome I with a region of 12.2-14.7 Mbp. This metabolite trait also had the highest heritability (H^2^ = 0.82), but without significant transgression.

To gain an overview of the genetic complexity and its effect on the detection of QTL, we identified the overlap between metabolite traits with significant mQTL, transgressive segregation, and heritability (Figure 5B). Interestingly, 7/18 metabolite traits (39%) that display significant transgression also mapped to significant mQTL. 17/18 metabolite traits (94%) that have a significant transgression also had relatively high heritability. Furthermore, 25/51 metabolite traits (49%) that showed a moderate to high heritability were mapped to one or more significant mQTL. Overall, only seven metabolite traits with significant transgression and heritability were mapped to one or more QTL. These findings suggest that many metabolite traits showed heritable variations and the regulation of these metabolite traits are highly complex and likely involve interactions between different genetic variants. This is supported by an analysis of the heritable variation in the parental strains versus the H^2^ in the RILs (Supplementary file 5). Here we observed that most of the H^2^ is either driven by many additive loci with opposing effects or by genetic interactions such as epistasis.

### Independent confirmation of the C14:1 and C18:2 mQTL using introgression lines

Next, we decided to focus on two mQTL that displayed the most significant linkage and high heritability: the mQTL for C14:1 on chromosome I and that for C18:2 on chromosome IV. For C14:1, 34% of variation could be explained by the genotype at the peak location on chromosome I (Figure 6A-B), and RILs with a CB4856 genotype at this QTL were associated with higher levels of C14:1 than RILs with an N2 genotype at this QTL (Figure 6B). All these observations show that a major locus is affecting C14:1 levels between these strains.

**Figure 6.**
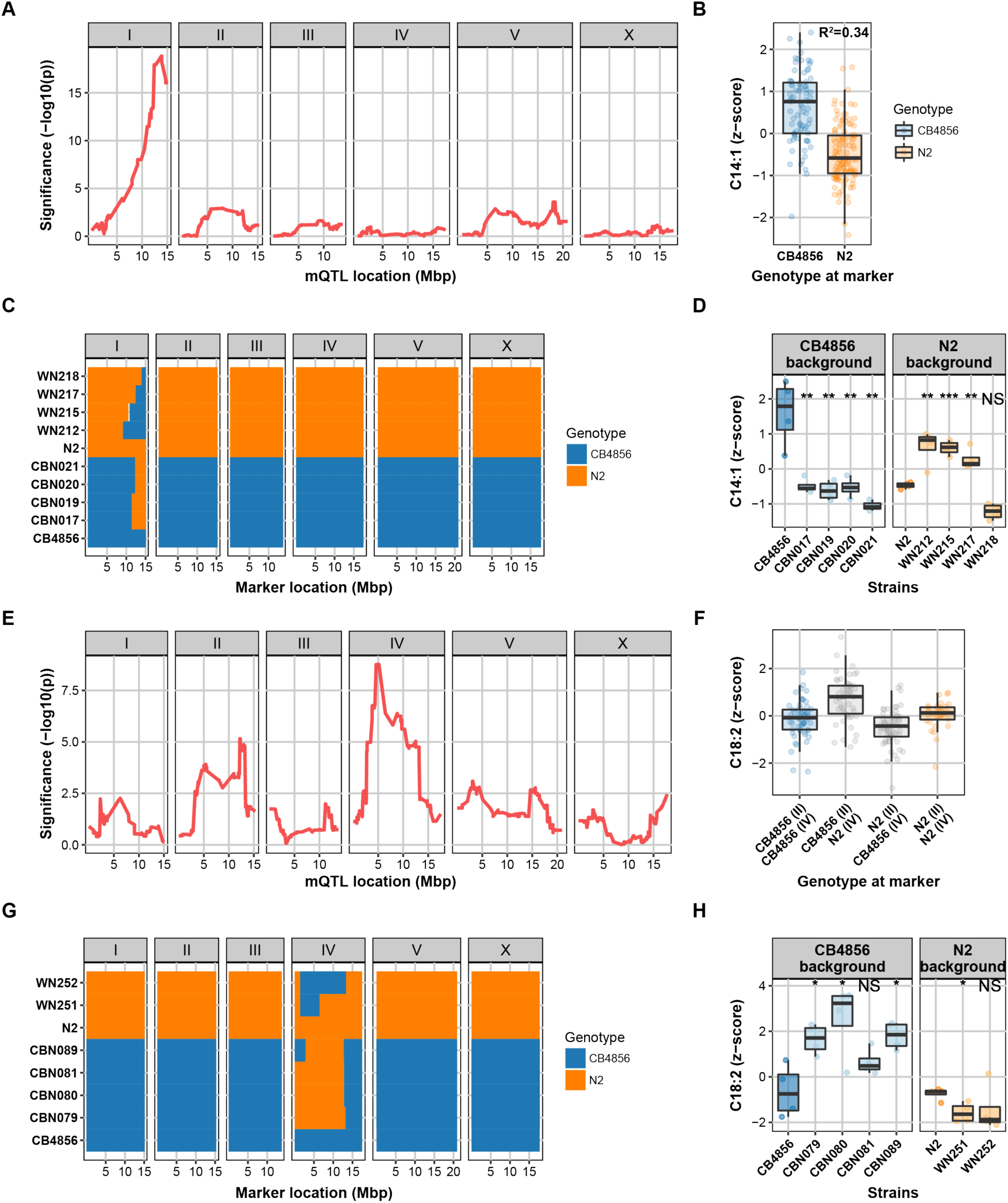
QTL peaks for C14:1 and C18:2 level and narrowing down the QTL peak regions with ILs, respectively. (A) A strong QTL for C14:1 was detected on chromosome I. (B) The genetic variations attribute to a QTL is calculated by the correlation between the metabolite level and the genotype at the peak location. RILs have a CB marker at this locus have a relatively higher abundance of C14:1 compared to those have an N2 marker. 34% of the variation in fatty acid C14:1 level can be explained by genetic variation on chromosome I. (C) Genotypes of the ILs with either a CB4856 background or an N2 background. The genome of an IL is composed of a recipient genome contributed by one of the parental strains and a short homozygous segment of the genome contributed by the other parental strain. WN212, WN215, WN217, and WN218 strains are ILs with an N2 background and CBN017, CBN019, CBN020, CBN021 are ILs with a CB4856 background. Genomic segments from N2 are marked with orange, those from CB4856 are marked with blue. (D) Metabolite profile of ILs and two parental strains. CB4856 has higher levels of C14:1. All CBN lines have lower levels than CB4856 (metabolite levels were normalized to percentage z-score), meaning that there is a QTL covered by all these lines, same for all WN lines, only WN218 has a lower level than N2. Therefore, this strain does not contain the QTL. The WN ILs narrow the QTL for C14:1 down to a region of 12.42-13.84 Mbp. Significance was calculated using Student’s *t*-test, and corrected for multiple testing. * *q*≤0.05; ** *q*≤0.01; *** *q*≤0.001, NS not significant. (E) Two QTL were detected for C18:2, one QTL peak was detected on chromosome II and one on chromosome IV. (F) RILs have an N2 marker at the locus on chromosome IV have a relatively higher abundance of C18:2 compared to those have an CB marker. 27% of the variation in fatty acid C18:2 level can be explained by genetic variation on chromosome II and IV. (G) Genotypes of the ILs with either a CB4856 background or an N2 background. CBN079, CBN080, CBN081, CBN089 are ILs with a CB4856 background and WN251, WN252 strains are ILs with an N2 background. Genomic segments from N2 are marked with orange, those from CB4856 are marked with blue. (H) Fatty acid profile of ILs and two parental strains. Three of the CBN lines have higher levels than CB4856 (metabolite levels were normalized to percentage z-score) and one of the two WN strains confirmed the mQTL for C18:2. The mQTL could be confirmed to a region of 2.8-6.4 Mbp. Significance was calculated using Student’s *t*-test, and corrected for multiple testing. * *q*≤0.05; ** *q*≤0.01; *** *q*≤0.001, NS not significant.

We next used introgression lines (ILs) to independently validate the C14:1 mQTL and narrow down the mQTL peak to a smaller region. These IL strains contain a small genomic segment derived from one parental strain introgressed in the genetic background of the other parental strain (Figure 6C). For C14:1, we used eight ILs (Sterken 2016). Four strains (WN212, WN215, WN217, and WN218) contain a CB4856-derived introgression in an N2 genetic background (Doroszuk et al. 2009), and four strains (CBN017, CBN019, CBN020, and CBN021) contain an N2-derived introgression in a CB4856 genetic background (Sterken 2016). The introgressions of both IL sets cover the C14:1 mQTL on chromosome I (Figure 6C). We measured the metabolite levels in the ILs and tested the hypotheses put forward from the RIL experiment (Figure 6D): (1) an N2 locus decreases the C14:1 abundance and (2) a CB4856 locus increases the abundance. We found that the four CB4856-background ILs had lower C14:1 levels, confirming the first hypothesis (FDR < 0.05). Three out of four N2-background ILs had increased C14:1 levels, which confirmed the second hypothesis (FDR < 0.05). Only WN218 did not show a significant increase in the C14:1 abundance (Figure 6D). By analyzing the introgression locations, we hence narrowed down the mQTL region to 12.4-13.8 Mbp.

For the FA C18:2, two QTL were identified, one on chromosome II (lower abundance in N2 genotype) and one on chromosome IV (higher abundance in N2 genotype) (Figure 6E-F). Together, these two loci explain 27% of the trait variation. For confirmation of the C18:2 mQTL, we used two N2 genetic background ILs (WN251 and WN252) and four CB4856 genetic background ILs (CBN079, CBN080, CBN081, and CBN089) (Figure 6G). As for C14:1, we tested the prediction from the IL panel: an increase in abundance by the CB4856 locus and a decrease by the N2 locus (in presence of the minor mQTL) (Figure 6H). Here, three of the four CB4856-background ILs and one of the two N2-background ILs confirmed the mQTL for C18:2 (FDR < 0.05). Based on the minimum region of the confirming ILs, the mQTL could be confirmed to the region spanning 2.8-6.4 Mbp.

Overall, these results independently confirmed the detection of the two mQTL in the RILs, demonstrating that this powerful approach is an important step to make forward genetics approaches possible in identifying complex genetics underlying metabolic regulation.

## Discussion

Imbalanced metabolic homeostasis often leads to chronic metabolic diseases (Gao et al. 2014). How genetic and environmental factors contribute to disturbed metabolism differs significantly among individuals and this process remains elusive (Andreux et al. 2012). Recent technological advances in “-omics” studies have made it possible to measure large-scale interactions in many components of a cell, allowing the study of complex biological systems and identification of new biochemical mechanisms (Williams et al. 2016). Of all - omics techniques, metabolomics most closely resembles the phenotype of a cell and is therefore an important area to study (Ramautar et al. 2013). Since mammalian model systems can be too complex to dissect the underlying mechanism of action in complex metabolic traits, we used the genetically tractable and cost-effective organism *C. elegans* in this study. To study the quantitative trait metabolite abundance, we measured FAs and AAs in a population of 199 *C. elegans* RILs derived from the genetically diverse strains N2 (Bristol) and CB4856 (Hawaii) (Li et al. 2006; Thompson et al. 2015; Gao et al. 2017a). This systems approach enabled us to investigate linkage between genetics and metabolism, identifying genomic regions contributing to genetic variation in metabolite abundances.

In hybrids of inbred lines and crosses between populations of diverse animals, transgressive or extreme phenotypes (transgressive segregation) falling beyond the parental phenotypes are often seen (deVicente and Tanksley 1993; Rieseberg et al. 1999). For example, in *C. elegans* transgression has been reported for some quantitative morphological traits, such as body size, egg size and fertility (Gutteling et al. 2007; Kammenga et al. 2007). For metabolite abundances, we found that 28.6% (18/63) of the metabolite traits displayed transgression. This hints that the genetic variation underlying metabolite abundance is a complex trait. Transgressive segregation in metabolite abundances is more extreme compared to transgression in gene-expression in *C. elegans* (affecting ~6% of the genes), (Vinuela et al. 2012). This makes sense in the light of metabolites as complex cellular phenotypes that can be affected on many levels.

We then calculated the broad-sense heritability to estimate the genetic contribution to the high levels of variation between the RILs we observed in the metabolite traits. The majority of metabolite traits were highly heritable, suggesting there was a high chance of identifying an associated genomic locus. Previous studies in RILs from other organisms have demonstrated that most metabolic traits are highly heritable (H^2^>0.5) (Andreux et al. 2012; Chen et al. 2014). Despite the fact that the heritability range differs across species, we showed ample evidence for moderate to high heritability in the metabolite traits of the 199 RILs with a range from 0.32 to 0.82, motivating us to identify detailed correlations between metabolites and the causal QTL for the metabolite traits.

Next, we analyzed the metabolite-metabolite correlations of the same metabolite classes and found strong correlations between multiple pairs of metabolite traits. Because involvement of different types of AAs in different metabolic pathways is rather complex, strong correlations between different pairs of AAs imply shared biological properties. All three BCAAs—valine, leucine, and isoleucine—showed strong positive correlation, likely because they share chemical similarities and are catabolized using similar pathways (Valerio et al. 2011; Mansfeld et al. 2015). In addition, two aromatic AAs tyrosine and phenylalanine were positively correlated together with all three BCAAs. Such correlation was previously observed in several type 2 diabetes studies in humans (Felig et al. 1969; Wang et al. 2011; Chen et al. 2016). For instance, in a nested case-control study for prediction of type 2 diabetes in a Framingham Offspring cohort, the abundance of these five AAs was significantly elevated under fasting state in high risk individuals (Wang et al. 2011). Our findings that these five AAs strongly correlate suggests a conserved regulatory role.

As metabolite traits are regulated in a complex fashion, QTL mapping allowed us to elucidate insights into the complexity of genetic regulation of metabolite abundances (Andreux et al. 2012). To increase the power of our approach, we chose to measure metabolite profiles in a large number of strains instead of multiple biological replicates per strain, as this has the highest impact on QTL mapping power (Andreux et al. 2012). Out of 36 significant mQTL that mapped for 26 metabolite traits, some metabolite traits were mapped to one or more mQTL, and some mQTL correlated with one or more metabolite traits. These findings indicate considerable genetic complexity in the regulation of metabolic quantitative traits (Flint and Mott 2001; Flint and Mackay 2009). To further investigate the genetic architecture of these quantitative phenotypes, we focused on the strongest mQTL detected for the FA C14:1 on chromosome I. Using ILs that covered the genomic region of this mQTL, we were able to narrow this mQTL down to a 1.4 Mbp region. In genetic studies with segregating populations, such as hybrid populations of mice and flies, large numbers of QTL could be detected, although most of them have very small effects and only few loci have moderate to large effects on quantitative traits (Flint and Mackay 2009). This C14:1 mQTL served as an excellent example as it showed a strong impact on the metabolite trait and explained large variations we have observed in the RILs, whereas the majority of the detected mQTLs only had a small effect.

Altogether, we believe that this RIL population of *C. elegans* provides us with a powerful platform to study the genetic basis of metabolism. The systems approach with QTL analysis makes it possible to address important questions related to genetic architectures of quantitative traits, such as genetic actions or interactions with each other and with the environment. The RILs can thereby play an important role to dissect the mechanisms underlying the complex processes of metabolism in a natural and unbiased manner and allow us to identify factors important for gene-by-environment interactions. In addition, this systems approach will also enable researchers to explore further into additional interventions, such as dietary alterations and environmental stresses-associated metabolic changes.

## Experimental procedures

### C. elegans strains and bacterial feeding strains

In total, 199 RILs were used (Li et al. 2006). ~25% of these RILs have been genotyped by sequencing (Thompson et al. 2015). A list of the strain names and their genotypes can be found in supplementary file (Supplementary file 1). For narrowing down the range of mQTL on chromosome I and IV 14 introgression lines (ILs) were used. Six of these ILs had an N2 genetic background: WN212, WN215, WN217, WN218, WN251, and WN252 (Doroszuk et al. 2009). Eight of these ILs had a CB4856 genetic background: CBN017, CBN019, CBN020, CBN021, CBN079, CBN080, CBN081, and CBN089 (Sterken 2016). The sequenced genotypes of WN212, WN217, WN251, and WN251 have been published in (Thompson et al. 2015), the sequenced genotypes of the strains WN218, and the CB4856 genetic-background strains are included in this manuscript (supplementary file 8).

### Strain culturing and experiments

Nematodes were cultured and maintained at 20°C on nematode growth media (NGM) agar plates. Culture conditions in all experiments were the same unless indicated otherwise. For metabolite profiling of 199 RIL strains, N2, and CB4856, age synchronized worms were obtained by alkaline hypochlorite treatment of gravid adults grown on *E. coli* OP50 lawn, 2000 eggs of each strain were then seeded onto NGM plates and cultured for 2.5 days allowing development to young adults. For heritability analysis of FAs and AAs, we collected worms in triplicates from 51 RIL strains that the genome composition contains high recombination (together with N2 and CB4856). To narrow down the QTL peak for C14:1 on Chromosome I and that for C18:2 on Chromosome IV, we prepared worm samples in four replicates for the IL strains.

### Whole genome sequence library prep and analysis for CB4856-background ILs

DNA was isolated from 100-300 µL of packed worms using Qiagen's Blood and Tissue kit (catalog # 69506). Following the ATL lysis step, 4 µL of 100 mg/mL RNAse was added to each sample and allowed to incubate for 2 min at room temperature. DNA concentration was determined using the Qubit dsDNA BR Assay Kit (catalog # Q32850). For each strain, a total of 0.75 ng of DNA was combined with 2.5 µL transposome (Illumina; kit # FC-121-1011) diluted 35x and 1X Tris Buffer (10X Tris Buffer: 100 mM Tris-HCl pH 8.0, 50 mM MgCl_2_) in a 10 µL final volume on ice. This reaction was incubated at 55°C for 10 min. The amplification reaction for each strain contained (final concentrations): 1X ExTaq Buffer, 0.2 mM dNTPs, 1 U ExTaq (Takara, catalog # RR001A), 0.2 µM primer 1, 0.2 µM primer 2, and 5 µL of tagmentation material from the previous step in a 25 µL total volume. Each strain had a unique pair of indexed primers. We first made a master mix containing buffer, water, dNTPs, and ExTaq then aliquot the appropriate volume of this mix into each well. We added the specific primer sets to each well and finally the tagmentation reaction. The amplification reaction was incubated in a thermocycler with the following conditions: 72°C for 3 min (1X); 95°C for 30 sec (1X); 95°C 10 sec, 62°C 30 sec, 72°C 3 min (20X); 10°C hold. We combined 8 µL from each amplification reaction to generate a pool of libraries. A portion of the libraries was electrophoresed on a 2% agarose gel. DNA was excised and gel purified using Qiagen's Gel Purification Kit (catalog # 28706). The libraries were sequenced on the Illumina HiSeq 2500 platform using a paired-end 100 bp reaction lane. Alignment, variant calling, and filtering were performed as previously (Cook et al. 2016). CB4856-background IL genotypes were called using the VCF file and a Hidden Markov Model as described previously (Cook and Andersen 2017).

### Metabolomics – Fatty acid extraction and MS analysis

Sample preparation for fatty acid (FA) extraction was followed as mentioned in our previous study (Gao et al. 2017a). A synchronized population of 2000 young adults was washed off the plates in M9 buffer and the worm pellet was washed with dH_2_O for three times and then collected in a 2 mL Eppendorf tube and freeze-dried overnight. Dried worm pellets were stored at room temperature until use. Dry worm pellets were re-suspended in 250 µL ice-cold 0.9% NaCl solution and homogenized with a 5 mm steel bead using a TissueLyser II (Qiagen) for 2x2.5 min at frequency of 30 times/sec, followed by a tip sonication (energy level: 40 joule; output: 8 watts) for two times on ice water. Protein quantification was performed with BCA assay.

Worm lysate (up to 150 µg protein) was transferred in a 4 mL FA-free glass vial, and 1 mL of freshly prepared 100% acetonitrile (ACN) / 37% hydrochloric acid (HCl) (4:1, v/v) was added to the lysate, together with deuterium-labeled internal standards. FA samples were hydrolyzed by incubating at 90°C for 2 h. After the vials cooled down to room temperature, 2 mL of hexane was added to the samples and mixed by vortexing for 5 sec followed by a centrifugation step at 1000 *g* for 1 min. The upper layer was transferred to an FA-free glass tube and evaporated at 30°C under a stream of nitrogen. FA residues were dissolved in 150 µL chloroform-methanol-water (50:45:5, v/v/v) solution containing 0.0025% aqueous ammonia, and then transferred to a Gilson vial for ESI-MS analysis.

### Metabolomics – Amino acid extraction and UPLC-MS/MS analysis

We used the same worm homogenate as mentioned and prepared for FA analysis. As described previously, amino acids (AAs) were extracted by transferring worm lysate (contains 50 µg of protein) to a 2 mL Eppendorf tube, and 1 mL of 80% ACN plus 20 µL of internal standard mixture were added to the lysate and homogenized by vortexing (Gao et al. 2017a). Samples were centrifuged and the supernatant was transferred to a 4 mL glass vial and evaporated under a stream of nitrogen at 40°C. After evaporation, AA residue was dissolved in 220 µl of 0.01% (v/v in MQ water) heptafluorobutyric acid. Then the suspension was transferred to a Gilson vial for HPLC-MS/MS analysis.

### Batch correction and data normalization

The metabolites were measured in five batches of FA and AA measurements, 400 samples in total. FA and AA measurements for N2 and CB4856 were repeated seven times, seven RILs were repeated five times, 44 RILs four times, 27 RILs two times, and 121 RILs one time (Supplementary file 2). Because of reliability detection limits of the MS platform, FA measurements with a concentration below 0.03 nmol/mg of protein were censored, as were AA concentrations below 0.4 nmol/mg of protein (Gao et al. 2017a). Since the measurements of the FA concentrations and the AA concentrations were conducted in the same sample, the amount measured was expressed as a ratio of the total composition. This was calculated independently for FAs and AAs using

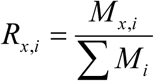

where R is the fraction of metabolite i (one of 40 FAs or one of 19 AAs) of all the metabolites (either FAs or AAs) measured in sample x. M is the concentration of metabolite i of sample x.

The fractions were batch corrected by

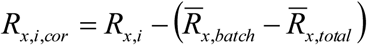

where R is the is the fraction of metabolite i in sample x, of which the difference between the batch average and the total average is subtracted.

Thereafter, the metabolite levels were expressed as a *z*-score by

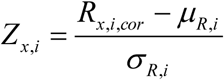

where Z is the *z*-score of metabolite i of sample x, and µ is the mean for that metabolite and σ is the standard deviation. This transformation was used in the further analysis. Outliers were removed if the trait value exceeded µ±2σ as measured per strain.

## Statistical analysis

### Software used

The data was analyzed in “R” (version 3.3.3, x64) using custom written scripts. The code required for analysis is made available via GitLab (https://git.wur.nl/mark_sterken/Metabolomics). In the analysis, the tidyverse package was used for organizing and plotting the data (www.tidyverse.org).

### Correlation analysis

The traits were analyzed for correlation by calculating the Pearson correlation between metabolites (FAs and AAs independently) on the metabolite levels.

### Transgressive segregation

Transgressive segregation was calculated as in (Brem and Kruglyak 2005). In short, to determine for which traits the RIL panel displayed transgressive segregation, we first calculated the mean trait value over all observations for the parental lines (separately for N2 and CB4856). Second, the standard deviation (σ) was calculated for the N2 and CB4856 observations separately, the pooled σ was used for transgression calculations. Transgressive segregation was calculated as the number of RILs of which the mean trait value exceeded μ ± 2*σ, where the parent with the lowest μ determines the low threshold and the parent with the highest μ determines the high threshold.

The significance of transgression was determined by permutation, where the trait values were randomized over the strain designations. Subsequently, the same test as described above was executed. The permutation was repeated 1000 times for each trait, where after the obtained values were used as the by-chance distribution; the FDR = 0.05 threshold was taken as the 50^th^ highest value.

### Heritability estimation

The heritability of the metabolite levels was calculated over a sub-set of RIL strains for which repeated measurements were conducted (51 RILs, n ≥ 3) and for metabolites that were consistently detected (> 100 observations). Using an ANOVA explaining the metabolite variation over the strains, the broad-sense heritability was calculated as

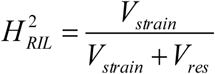

where H^2^ is the broad-sense heritability and V_strain_ is the variation explained by strain and V_res_ is the residual variation. The significance of the heritability was calculated by permutation, where the trait values were randomly assigned to strains. Over these permutated values, the variance captured by strain and the residual variance were calculated. This procedure was repeated 1000 times for each trait. The obtained values were used as the by-chance distribution and an FDR = 0.05 was taken as the 50^th^ highest value.

In the parental strains (n = 11 for both N2 and CB4856) the heritability was calculated by ANOVA, using

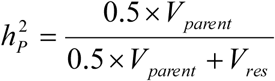

where h^2^ is the heritability, and V_parent_ is the variation explained by the parental genotypes and V_res_ is the residual variation. The factor 0.5 corrects for the overestimation of the additive variation in inbred strains (Hegmann and Possidente 1981). The same permutation approach as for the broad-sense heritability was applied, taking the FDR = 0.05 threshold as significant.

### Quantitative trait locus mapping

Quantitative trait loci (QTL) were mapped using custom scripts in “R”. The metabolite levels were fitted in a single marker model,

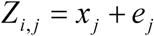

where Z is the *z*-score averaged over all strain replicates for metabolite i (one of 39 FAs or one of 19 AAs that were reliably measured in >100 strains) of RIL j (n = 199). This is explained over the genotype (either CB4856 or N2) on marker location x (x = 1, 2, …, 729) of RIL j.

### QTL threshold determination & power analysis

In order to account for multiple testing, the genome-wide significance threshold was determined via permutation. Here the *z*-scores per metabolite were randomly distributed over the genotypes. This permutated dataset was thereafter used in the same single marker model for QTL-mapping. This procedure was repeated for 100 randomized datasets. From these randomized mappings, a false discovery rate was determined based on multiple testing under dependency (Benjamini and Yekutieli 2001),

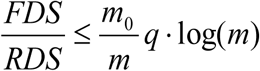

where false discovery (FDS) is the outcome of the permutations and real discovery (RDS) is the outcome of the expression QTL (eQTL) mapping at a specific significance level. The value of m_0_, the number of true null hypotheses tested, was 56-RDS and for the value of m, the number of hypotheses tested, the number of metabolites (56) was taken. The q-value was set at 0.05. In this way, a –log10(p) > 3.7 was found. The eQTL confidence interval was determined by a 1.5 drop in the –log10(p) value as measured from the eQTL peak.

The statistical power at the FDR threshold was determined by simulation. Using the genetic map of the 199 strains used in this study, QTL were simulated for each marker location. Per location 10 QTL were simulated, explaining 5-80% of the variation (in increments of 5%). In order to simulate technical noise, we introduced random variation based on a standard normal distribution (σ = 1, μ = 0). The simulated peak size was set correspondingly (*e.g.* 50% explained variation corresponds to a peak size of 2 in the simulated noise). Based on the set permutation threshold (–log10(p) > 3.7), the number of correctly detected QTL, the number of false-positives, and the number of undetected QTL were counted. From the simulation, we also inferred the precision of the mapping, by evaluating the effect size estimation and the QTL location (based on a –log10(p) drop of 1.5). The detailed results of the analysis can be found in Supplementary file 6.

### Introgression line analysis

The transformed metabolomics data on C14:1 and C18:2, obtained in 8 ILs covering the C14:1 mQTL and 6 ILs covering the C18:2 mQTL, was compared to the genetic background parent. This means that the ILs with the CB4856 genetic background were compared to CB4856 and the ILs with the N2 genetic background were compared to N2. For both loci separately, we tested the differences in C14:1 and C18:2 abundance using a Student’s t-test, explicitly testing the hypothesis formed from the RIL data. Therefore, for C14:1 the CB4856-background ILs were tested for lower trait levels compared to the genetic background strain and the N2-background ILs were tested for higher trait levels compared to the genetic background strain. For C18:2 the CB4856-background ILs were tested for higher trait levels compared to the genetic background strain and the N2-background ILs were tested for lower trait levels compared to the genetic background strain. The significances were adjusted for multiple testing using the Benjamini & Hochberg correction, as implemented in the p.adjust function in “R” (Benjamini and Hochberg 1995).

## Acknowledgements

AWG is supported by an AMC PhD Scholarship. Work in the Houtkooper group is financially supported by an ERC Starting grant (no. 638290), and a VIDI grant from ZonMw (no. 91715305). The authors thank Myrthe Walhout and Joost Riksen for helping with constructing the CBN introgression lines and Daniel Cook, Robyn Tanny, and Erik Andersen for help in sequencing these strains.

## Author contributions

AWG, MGS, JEK, and RHH conceived and designed the project. AWG, JB, RK, LBS, and MGS performed experiments and interpreted data, JB and MGS performed bioinformatics. JvC and MGS constructed CB4856 genetic background ILs. AWG, MGS, JB, JEK, and RHH wrote the manuscript, with contributions from all other authors.

## Competing financial interests

The authors declare no competing financial interests related to this work.

## Additional Information Supplementary information

**Supplementary file 1:** A matrix of the genetic map of the strains used in this study. The first column contains the marker names, followed by a column with the chromosome and a column with the base-pair position (WS258). The following columns contain the strains used in this study. A ‘1’ denotes an N2 genotype, a ‘-1’ a CB4856 genotype.

**Supplementary file 2**: A table with the metabolomics data gathered for this study. The file includes the raw values as obtained via the MS analysis and all the data transformations used for subsequent analysis. Data is ordered by batch, metabolite, strain, and biological replicate. For each trait, an upper and lower bound for outlier analysis is indicated.

**Supplementary file 3**: The outcome of the transgression analysis. Per trait it is indicated how many RILs fell beyond the mean ± pooled standard deviation of the parental strains N2 and CB4856 (n_strains_transgression). The FDR0.1, FDR0.05, and FDR0.01 give the number of strains that constitute the significance boundary as determined by permutation at a false discovery rate of 0.1, 0.05, and 0.01 respectively.

**Supplementary file 4**: The outcome of the heritability analysis. For both the parental strains N2 and CB4856 (H2_ANOVA_PL) and the RIL population (H2_ANOVA) the heritability is given per metabolite. The individual components Vg (genetic variance) and Ve (environmental variance) are also given, as is the false discovery rate threshold of 0.05 (FDR0.05) for both populations. The significance column indicates which populations displayed significant heritability.

**Supplementary file 5**: A plot of the heritability calculated in the parental strains versus the broad sense heritability calculated in the RILs. The colors indicate the significance of the heritability estimate (as determined by permutation, false discovery rate < 0.05). If the broad-sense heritability exceeds the heritability in the parental strains, it indicates a complex trait architecture (multiple regulatory loci).

**Supplementary file 6**: Analysis of the statistical power of the QTL mapping. At each marker location 10 peaks were simulated that explained 5-80% of the variation (in 5% increments). For each simulated peak, it was determined if the QTL was detected at the right location. Additionally, the ratio of the estimated effect size versus the simulated effect size is given for the simulations (in quantiles), as is the precision of the QTL location (in quantiles).

**Supplementary file 7**: A summary table of the mQTL. Per metabolite the mQTL are given, the significance of the mQTL in –log10(p), the effect of the mQTL (positive numbers indicate higher at N2 loci, negative numbers indicate higher at CB4856 loci), and the amount of variation the mQTL explains (according to a single marker model). Also, the mQTL location is given: chromosome (qtl_chromosome), peak location in base pairs (qtl_bp) and in marker location (qtl_marker_peak). The confidence interval of the mQTL location, based on a 1.5 drop in –log10(p), is given (the left and right locations). Finally, the estimated heritabilities are included per metabolite.

**Supplementary file 8**: The genotypes of the WN218 strain and the eight CB4856-genetic background introgression lines used in this study. For the introgression regions, the number of supportive genetic variants over the total genetic variants is given (as determined by the hidden markov model).

---

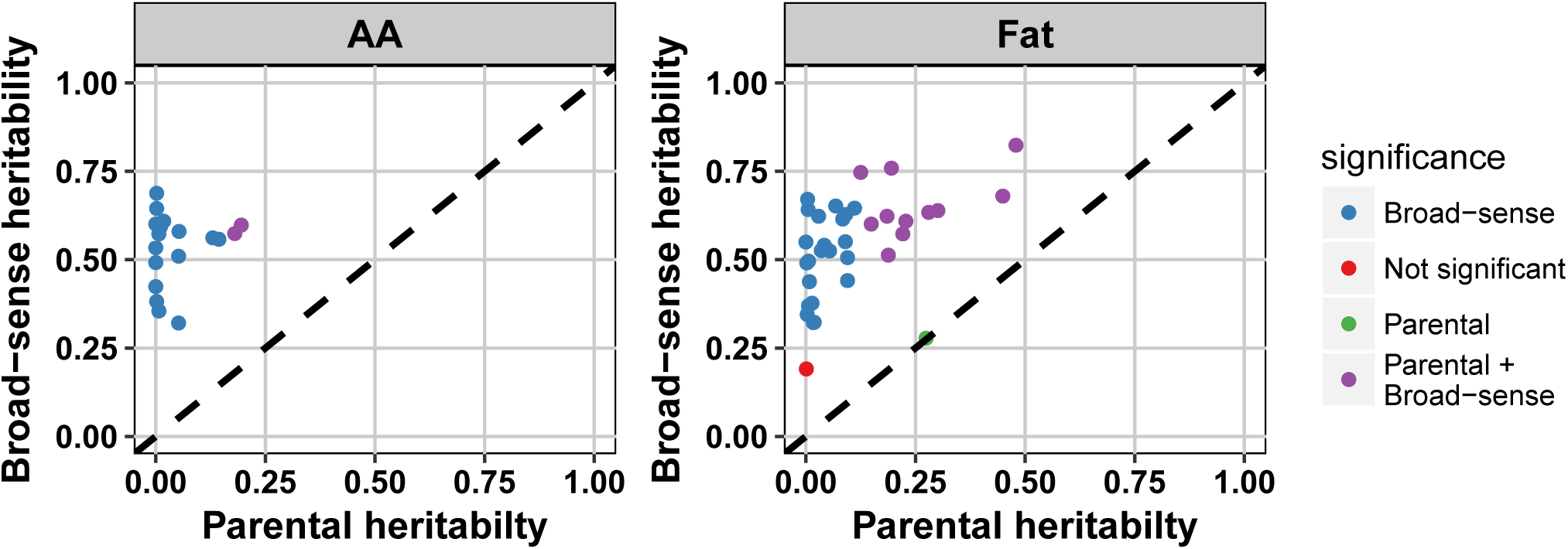

## References

Andersen EC, Gerke JP, Shapiro JA, Crissman JR, Ghosh R, Bloom JS, Felix. MA, Kruglyak L. 2012. Chromosome-scale selective sweeps shape Caenorhabditis elegans genomic diversity. Nat Genet 44: 285–290.

Andreux PA, Williams EG, Koutnikova H, Houtkooper RH, Champy MF, Henry H, Schoonjans K, Williams RW, Auwerx J. 2012. Systems genetics of metabolism: the use of the BXD murine reference panel for multiscalar integration of traits. Cell 150: 1287–1299.

Benjamini Y, Hochberg Y. 1995. Controlling the False Discovery Rate: A Practical and Powerful Approach to Multiple Testing. Journal of the Royal Statistical Society Series B (Methodological) 57: 289–300.

Benjamini Y, Yekutieli D. 2001. The control of the false discovery rate in multiple testing under dependency. The Annals of Statistics 29: 1165–1188.

Brem RB, Kruglyak L. 2005. The landscape of genetic complexity across 5,700 gene expression traits in yeast. Proceedings of the National Academy of Sciences of the United States of America 102: 1572–1577.

Chen T, Ni Y, Ma X, Bao Y, Liu J, Huang F, Hu C, Xie G, Zhao A, Jia W et al. 2016. Branched-chain and aromatic amino acid profiles and diabetes risk in Chinese populations. Scientific reports 6: 20594.

Chen W, Gao Y, Xie W, Gong L, Lu K, Wang W, Li Y, Liu X, Zhang H, Dong H et al. 2014. Genome-wide association analyses provide genetic and biochemical insights into natural variation in rice metabolism. Nat Genet 46: 714–721.

Cook DE, Andersen EC 2017. VCF-kit: assorted utilities for the variant call format. Bioinformatics 33: 1581–1582.

Cook DE, Zdraljevic S, Tanny RE, Seo B, Riccardi DD, Noble LM, Rockman MV, Alkema MJ, Braendle C, Kammenga JE et al. 2016. The Genetic Basis of Natural Variation in Caenorhabditis elegans Telomere Length. Genetics 204: 371–383.

deVicente MC, Tanksley SD. 1993. QTL analysis of transgressive segregation in an interspecific tomato cross. Genetics 134: 585–596.

Doroszuk A, Snoek LB, Fradin E, Riksen J, Kammenga J. 2009. A genome-wide library of CB4856/N2 introgression lines of Caenorhabditis elegans. Nucleic Acids Res 37: e110.

Felig P, Marliss E, Cahill GF Jr. 1969. Plasma amino acid levels and insulin secretion in obesity. The New England journal of medicine 281: 811–816.

Flint J, Mackay TF. 2009. Genetic architecture of quantitative traits in mice, flies, and humans. Genome Res 19: 723–733.

Flint J, Mott R. 2001. Finding the molecular basis of quantitative traits: successes and pitfalls. Nat Rev Genet 2: 437–445.

Gao AW, Canto C, Houtkooper RH. 2014. Mitochondrial response to nutrient availability and its role in metabolic disease. EMBO molecular medicine doi:10.1002/emmm.201303782.

Gao AW, Chatzispyrou IA, Kamble R, Liu YJ, Herzog K, Smith RL, van Lenthe H, Vervaart MAT., van Cruchten A, Luyf AC et al. 2017a. A sensitive mass spectrometry platform identifies metabolic changes of life history traits in C. elegans. Scientific reports 7: 2408.

Gao AW, Uit de Bos J., Sterken MG, Kammenga JE, Smith RL, Houtkooper RH. 2017b. Forward and reverse genetics approaches to uncover metabolic aging pathways in Caenorhabditis elegans. Biochimica et biophysica acta doi:10.1016/j.bbadis.2017.09.006.

Gutteling EW, Riksen JA, Bakker J, Kammenga JE. 2007. Mapping phenotypic plasticity and genotype-environment interactions affecting life-history traits in Caenorhabditis elegans. Heredity (Edinb) 98: 28–37.

Hegmann JP, Possidente B. 1981. Estimating genetic correlations from inbred strains. Behav Genet 11: 103–114.

Houtkooper RH, Mouchiroud L, Ryu D, Moullan N, Katsyuba E, Knott G, Williams RW, Auwerx J. 2013. Mitonuclear protein imbalance as a conserved longevity mechanism. Nature 497: 451–457.

Houtkooper RH, Williams RW, Auwerx J. 2010. Metabolic Networks of Longevity. Cell 142: 9–14.

Kammenga JE, Doroszuk A, Riksen JA, Hazendonk E, Spiridon L, Petrescu AJ, Tijsterman M, Plasterk RH, Bakker J.. 2007. A Caenorhabditis elegans wild type defies the temperature-size rule owing to a single nucleotide polymorphism in tra-3. PLoS Genet 3: e34.

Li Y, Alvarez OA, Gutteling EW, Tijsterman M, Fu J, Riksen JA, Hazendonk E, Prins P, Plasterk RH, Jansen RC et al. 2006. Mapping determinants of gene expression plasticity by genetical genomics in C. elegans. PLoS Genet 2: e222.

Lopez-Otin C, Blasco MA, Partridge L, Serrano M, Kroemer G. 2013. The hallmarks of aging. Cell 153: 1194–1217.

Mansfeld J, Urban N, Priebe S, Groth M, Frahm C, Hartmann N, Gebauer J, Ravichandran M, Dommaschk. A, Schmeisser S et al. 2015. Branched-chain amino acid catabolism is a conserved regulator of physiological ageing. Nat Commun 6: 10043.

Ramautar R, Berger R, van der Greef J, Hankemeier T. 2013. Human metabolomics: strategies to understand biology. Curr Opin Chem Biol 17: 841–846.

Rieseberg LH, Archer MA, Wayne RK. 1999. Transgressive segregation, adaptation and speciation. Heredity (Edinb) 83 (Pt 4): 363–372.

Rockman MV, Skrovanek SS, Kruglyak L. 2010. Selection at linked sites shapes heritable phenotypic variation in C. elegans. Science 330: 372–376.

Rodriguez M, Snoek LB, Riksen JA, Bevers RP, Kammenga JE. 2012. Genetic variation for stress-response hormesis in C. elegans lifespan. Exp Gerontol 47: 581–587.

Slate J. 2005. Quantitative trait locus mapping in natural populations: progress, caveats and future directions. Mol Ecol 14: 363–379.

Snoek BL, Sterken MG, Bevers RPJ, Volkers RJM, Van’t Hof A, Brenchley R, Riksen JAG, Cossins A, Kammenga JE. 2017. Contribution of trans regulatory eQTL to cryptic genetic variation in C. elegans. BMC Genomics 18: 500.

Sterken MG. 2016. Building towards a multi-dimensional genetic architecture in Caenorhabditis elegans, doi:10.18174/386549, p. 168. Wageningen University.

Sterken MG, Snoek LB, Kammenga JE, Andersen EC. 2015. The laboratory domestication of Caenorhabditis elegans. Trends Genet 31: 224–231.

Thompson OA, Snoek LB, Nijveen H, Sterken MG, Volkers RJ, Brenchley R, Van’t Hof A, Bevers RP, Cossins AR, Yanai I et al. 2015. Remarkably Divergent Regions Punctuate the Genome Assembly of the Caenorhabditis elegans Hawaiian Strain CB4856. Genetics 200: 975–989.

Valerio A, D’Antona G, Nisoli E. 2011. Branched-chain amino acids, mitochondrial biogenesis, and healthspan: an evolutionary perspective. Aging 3: 464–478.

Vinuela A, Snoek LB, Riksen JA, Kammenga JE. 2010. Genome-wide gene expression regulation as a function of genotype and age in C. elegans. Genome Res 20: 929–937.

Vinuela A, Snoek LB, Riksen JA, Kammenga JE. 2012. Aging Uncouples Heritability and Expression-QTL in Caenorhabditis elegans. G3 (Bethesda) 2: 597–605.

Visscher PM, Wray NR, Zhang Q, Sklar P, McCarthy MI, Brown MA, Yang J. 2017. 10 Years of GWAS Discovery: Biology, Function, and Translation. American journal of human genetics 101: 5–22.

Wang TJ, Larson MG, Vasan RS, Cheng S, Rhee EP, McCabe E, Lewis GD, Fox CS, Jacques PF, Fernandez C et al. 2011. Metabolite profiles and the risk of developing diabetes. Nature medicine 17: 448–453.

Williams EG, Auwerx J. 2015. The Convergence of Systems and Reductionist Approaches in Complex Trait Analysis. Cell 162: 23–32.

Williams EG, Wu Y, Jha P, Dubuis S, Blattmann P, Argmann CA, Houten SM, Amariuta T, Wolski W, Zamboni N et al. 2016. Systems proteomics of liver mitochondria function. Science 352: aad0189.

